# Accelerating surveillance and research of antimicrobial resistance – an online repository for sharing of antimicrobial susceptibility data associated with whole genome sequences

**DOI:** 10.1101/532267

**Authors:** Sébastien Matamoros, Rene. S. Hendriksen, Balint Pataki, Nima Pakseresht, Marc Rossello, Nicole Silvester, Clara Amid, COMPARE ML- AMR group, Guy Cochrane, Istvan Csabai, Ole Lund, Constance Schultsz

**Affiliations:** Amsterdam UMC, University of Amsterdam, Department of Medical Microbiology, Amsterdam, The Netherlands.; National Food Institute, Technical University of Denmark, Lyngby, Denmark.; Department of Physics of Complex Systems, ELTE Eötvös Loránd University, Budapest, Hungary.; Department of Computational Sciences, Wigner Research Centre for Physics of the HAS, Budapest, Hungary.; European Molecular Biology Laboratory (EMBL), European Bioinformatics Institute (EBI), Wellcome Genome Campus, Hinxton, Cambridge, CB10 1SD, UK.; Amsterdam UMC, University of Amsterdam, Department of Global Health, Amsterdam Institute for Global Health and Development, Amsterdam, The Netherlands.; Fondation Mérieux, Lyon, France.; Department of Infectious Diseases, Robert Koch Institut, Berlin, Germany.; Department of Medical Microbiology, Vaccine & Infectious Disease Institute, Antwerp University, University Hospital Antwerp, Antwerp, Belgium.; Department of Physics and Astronomy (DIFA), University of Bologna, Bologna, Italy.; Department of Agricultural and Food Sciences (DISTAL), University of Bologna, Bologna, Italy.; Animal and Plant Health Agency, Addlestone, Surrey,United Kingdom.; Statens Serum Institut, Denmark.; Oxford University Clinical Research Unit, Centre for Tropical Medicine, Ho Chi Minh City, Vietnam.

## Abstract

Antimicrobial resistance (AMR) is an emerging threat to modern medicine. Improved diagnostics and surveillance of resistant bacteria require the development of next generation analysis tools and collaboration between international partners. Here, we present the “AMR data hub”, an online infrastructure for storage and sharing of structured phenotypic AMR data linked to bacterial genome sequences.

Leveraging infrastructure built by the European COMPARE Consortium and structured around the European Nucleotide Archive (ENA), the AMR data hub already provides an extensive data collection for some 500 isolates with linked genome and AMR data. Representing these data in standardized formats, we provide tools for the validation and submission of new data and services supporting search, browse and retrieval.

The current collection was created through a collaboration by several partners from the European COMPARE Consortium, demonstrating the capacities and utility of the AMR data hub and its associated tools. We anticipate growth of content and offer the hub as a basis for future research into methods to explore and predict AMR.

## Introduction

Antimicrobials are widely regarded as one of the major advances in modern medicine ^1^. The global emergence, however, of antimicrobial resistance (AMR) threatens the very core of modern medicine with the potential to turn the global population back in time to the pre-antibiotic era in which simple surgical procedures and common infections could have deadly consequences ^2^.

The decreased costs of next generation sequencing (NGS) combined with the progress made in big data analysis such as machine leaning (ML) represent innovative opportunities to tackle the AMR crisis^3^. Many bacterial phenotypic traits, including AMR, can be directly linked to the presence of genomic determinants such as genes, Single Nucleotide Polymorphism’s (SNPs) or transcription promoters which can be identified using functional genomics approaches on large databases of genomes. Recent studies have used computational approaches such as ML to predict antimicrobial susceptibility from genomic data or to discover previously unidentified antibiotic resistance determinants^4–6^. Today, the major limitation for such approaches is not the lack of advanced computational methods or hardware resources but the lack of large enough well curated, annotated data sets where phenotypic AMR data and genomes are linked.

Academic research initiatives and public health organisations could benefit from the implementation of online repositories capable of storing large amounts of genome sequence and antimicrobial susceptibility testing (AST) data. ^7^. For example, the PATRIC database (https://patricbrc.org/ ^6^) has been used for development of AMR prediction algorithms, but is a closed architecture using a project specific data entry template. The NCBI is offering a similar service, linking antibiograms with genomes deposited in the Sequence Repository Archive (SRA), but all data have to be made public immediately (https://www.ncbi.nlm.nih.gov/pathogens/isolates#/search/).

Different stakeholders in the AMR field may have different requirements regarding the accessibility of AST data. Making optimal use of the opportunities described above requires enabling the global sharing of data, but some institutions are reluctant to immediately make their AST data publicly accessible for privacy, legal or other reasons ^8^. National public health institutes would be encouraged to create supra-national networks using a standard format to share data for AST result analysis, while academics would find a solution to share post-publication data, encouraging reproducibility and cross-validation experiments, if such a database structure would become available. Thus, we have identified a clear need for a database structure that can support public AST data as well as those data that are to be shared privately for a period of time until publication.

The Horizon 2020 funded EU consortium COMPARE aims at bringing NGS to public health and clinical practice (http://www.compare-europe.eu/). European experts in AMR working within the COMPARE consortium, including the European Bioinformatics Institute (EMBL-EBI), which is part of the International Nucleotide Sequence Database Collaboration (INSDC; http://www.insdc.org/), have deployed the “data hub” system to allow sharing of isolate NGS and linked phenotypic AMR data. The data hub system allows data providers, such as public health and clinical laboratories, food safety agencies and veterinary institutes, to share and download genomic and related data sets. Data can either be kept private (pre-publication) or released as open-access at the discretion of the data providers (Amid *et al.*, 2019, *in preparation*). Novel software was developed for use in the AMR data hub that validates the conformity of submitted datasets. The system supports both qualitative and quantitative AST data such as those resulting from disk diffusion and micro-broth dilution tests. The AMR data hub “Notebook” reporting system has been configured to support rapid data mining of content.

Here, we present the AMR data hub, which permits sharing of large amounts of information that could be used for ML and other data analysis approaches, eventually resulting in accurate and quantitative, hence clinically relevant, predictions of AMR phenotypes based on NGS data.

## The AMR data hub

### The data hub system

The data hub system has been built as a broad infrastructure for the sharing and analysis of pathogen NGS data and related data types. Here is presented a brief outline of the system while a full description is provided in Amid *et al.*, (2019, in preparation). The data hubs are provided upon the foundation of the European Nucleotide Archive (ENA; https://www.ebi.ac.uk/ena), an open repository for sequence and related data ^9^. The concept was developed and introduced as a model for rapid sharing of data and analysis outputs in public and pre-publication confidential status within the COMPARE consortium. Data hubs are restricted (by login and password) to the members of the project authorized to access the data. Pre-publication confidential data sharing between partners has been considered only in projects where immediate release of data/metadata was not possible, i.e. sensitive content has been awaiting publication, but partners needed access to confidential data for analysis. Ultimately, all data archived in data hubs are released into the public domain (the standard ENA database) after a period defined by data owners. Data and metadata reported by data providers are submitted to the hubs through systematic processes supported by a number of tools. Subsequently, structured and accessioned data/metadata are available for sharing between data consumers who have received consent from data providers. Data appropriate for a given computational analysis are selected and fed autonomously through cloud-based analysis workflows of which the outputs – “derived” data products – are fed back into the system.

### The antibiogram

In order to represent AST data, we have defined a new data type, the “antibiogram”, for use within the AMR data hub and, more broadly, within ENA. This new data type leverages the extensible “analysis object” system, with the addition of a new class specifically for the storage of phenotypic AMR data, designated “AMR_ANTIBIOGRAM”. Antibiograms are treated as data objects within the system and are supported in data submission and access services. As a new data type, building this support has required the development of open software that is distributed publicly (https://github.com/EBI-COMMUNITY/compare-amr) and used internally at EMBL-EBI for the validation and submission of incoming AST data.

We have aimed with the antibiogram for a format that is flexible and complete. Minimum requirements include, for each combination of isolate/ antibiotic provided, INSDC Sample accession (SAM*); species; antibiotic name; antibiotic susceptibility testing standard; breakpoint version; antibiotic susceptibility test method; measurement; measurement units; measurement sign; susceptibility phenotype; and test platform. Any combination of bacterial species and antimicrobial is supported. Reported antimicrobial susceptibility can be measured by microbroth dilution or zone diameter, all major testing platforms (Sensititre, VITEK and Phoenix) and standards (EUCAST or CLSI) are accepted. More uncommon methods can be added by using the free-text format of these sections. To help data providers, a detailed protocol explaining the preparation of the metadata form is presented on the GitHub repository of the project (https://github.com/EBI-COMMUNITY/compare-amr) along with tools for batch creation of antibiograms from excel files. An interactive web page allowing the manual creation of antibiograms is in preparation. It will provide an easy alternative for data providers registering a limited number of samples. Finally, a tutorial explaining the steps required to retrieve data (genomes and antibiograms) from the datahub is available on the GitHub repository.

Each antibiogram is linked to a bacterial genome within ENA. The association is asserted by linking the analysis object, i.e. the antibiogram, with the corresponding study, example: https://www.ebi.ac.uk/ena/data/view/PRJEB14981. As with other data deposited in the ENA, antibiograms can be kept confidential for a provider-defined period but must ultimately be released into public view. Antibiograms can be queried and retrieved through the AMR data hub (while confidential) as well as (when made public) through other alternatives, such as the Pathogen data portal (https://www.ebi.ac.uk/ena/pathogens/home^1^), a Discovery Application Programming Interface (API - https://www.ebi.ac.uk/ena/portal/api/), the ENA browser (https://www.ebi.ac.uk/ena) and services providing high-volume data access such as the ENA File Downloader (https://github.com/enasequence/ena-ftp-downloader/) for public data. Using the Pathogen data portal or the API, various filters can be used to refine the query such as bacterial species, country of origin, host, and more.

### Visualisation tools

To visualize the contents of the AMR data hub, a Notebook was configured (Figure 1) that has several options for comparison of a defined set of parameters from the database, such as the distribution of the minimal inhibitory concentration (MIC) of different antimicrobials as a function of the country of origin, or the comparison of MIC distributions between different antimicrobials. As such, this functionality can be used for surveillance purposes, providing a rapid overview of MIC distribution for a specific collection of isolates and how this compares to isolates from the same host or from different geographical regions.

**Figure 1:**
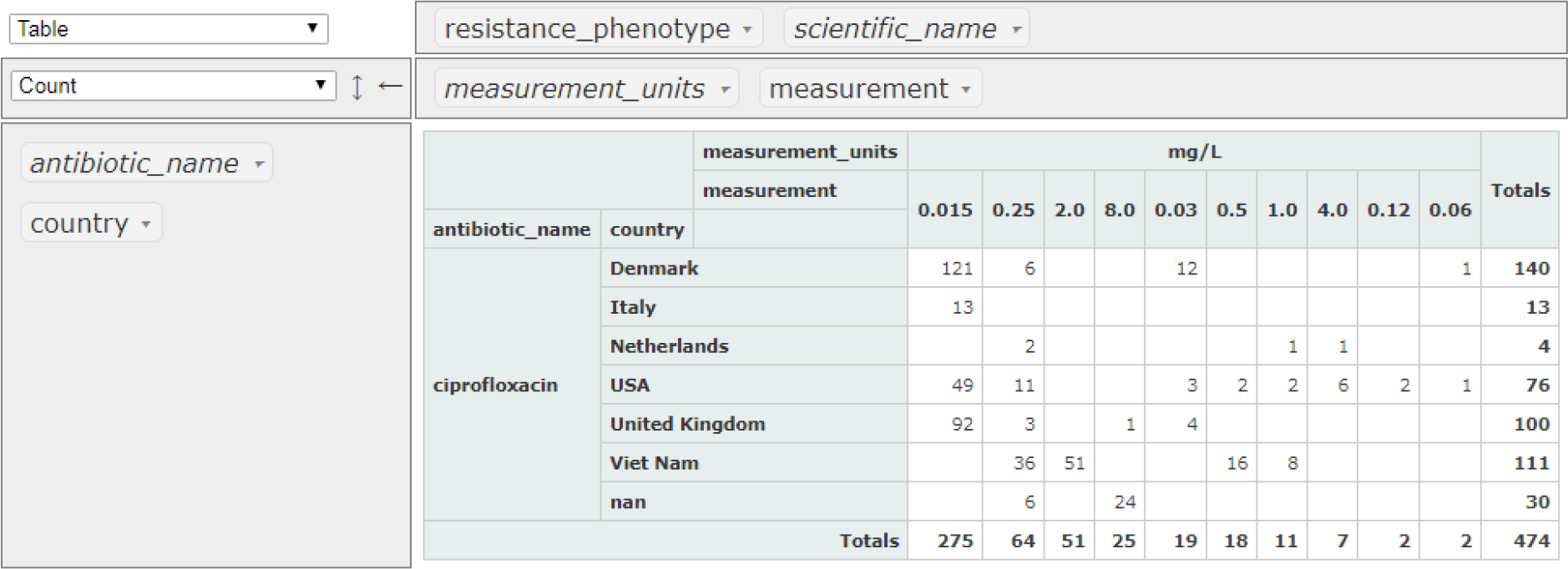
Visualization of the AST data deposited in the database on 15/10/2018. Data were filtered for: ciprofloxacin (antibiotic_name); *E. coli* (scientific_name) and mg/L (measurement_units).

The Notebook is integrated into the Pathogen portal and can be accessed from https://www.ebi.ac.uk/ena/pathogens/home under the “Explore” tab. Access to pre-publication data in the AMR data hub requires authentication via login and password with authorization to the corresponding project.

## Current content

### Statistics

As of 15-10-2018 the AMR data hub contains 577 *Escherichia coli* genomes with attached AST data, 1,842 *Salmonella enterica* and 16 *Enterococcus faecium* originating from 9 different countries. Data on susceptibility against 55 different antibiotics (or combinations) have been entered in the database so far. As an example, 577 *E. coli* isolates originating from seven different countries (Bangladesh; Denmark; Italy; Netherlands; UK; USA and Vietnam) have been tested for ciprofloxacin susceptibility, 474 by various dilution-based methods and 103 by disk diffusion.

A total of 470 *E. coli* antibiograms were submitted directly by COMPARE partners while 107 were imported from the US CDC database (n = 31) or previous publications (n = 76) ^10,11^.

## Distribution of the MICs

As shown in Figure 2, the distribution of the *E. coli* ciprofloxacin MICs, as an example, recorded in the present database follows a similar pattern as the ciprofloxacin MICs distribution reported by EUCAST which is based on more than 20,000 isolates (https://mic.eucast.org/Eucast2/).

**Figure 2:**
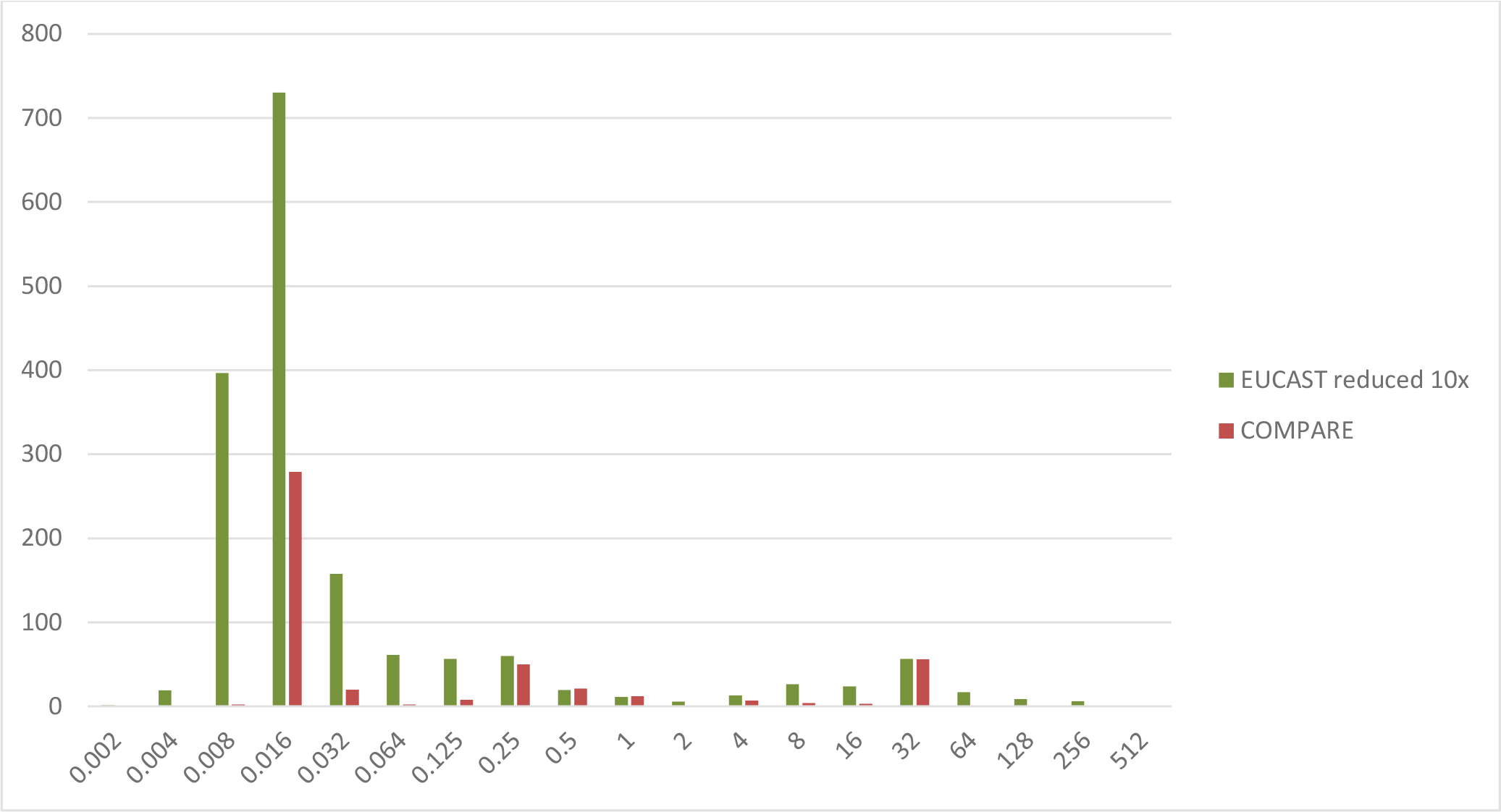
Distribution of the *E. coli* ciprofloxacin MICs and comparison with ECOFFs (https://mic.eucast.org/Eucast2/).

### Phylogenetic analysis

As an example of the possibilities offered by the AMR data hub, the entire collection of *E. coli* genomes was downloaded and a phylogenetic analysis of the population, in relation to ciprofloxacin MICs, was performed (methods in supplementary materials). As shown in Figure 3, this population is highly diverse and features a large range of ciprofloxacin MICs. Several isolates tend to cluster by country, such as those from Denmark or Vietnam. Additionally, a strong association between resistance and country can be observed. Most isolates from Vietnam show an MIC higher than 2 mg/ml. The study during which these isolates were collected focused on the presence of ESBL genes in *E. coli* ^12^ and it is possible that ciprofloxacin resistance was co-selected for, as was previously suggested ^13^. Additionally, 24 out of 30 isolates retrieved from the CDC antimicrobial resistance isolate bank (https://www.cdc.gov/drugresistance/resistance-bank/currently-available.html) exhibited an MIC of 8 mg/ml. This unusually high proportion of resistant isolates can be explained by the purpose of the CDC database, which is to provide a panel of well characterized resistant bacteria for testing of diagnostic devices and new antibiotic agents. Conversely, isolates from Denmark were collected as part of a routine surveillance effort from the veterinary institute and appear to all be susceptible to ciprofloxacin.

**Figure 3:**
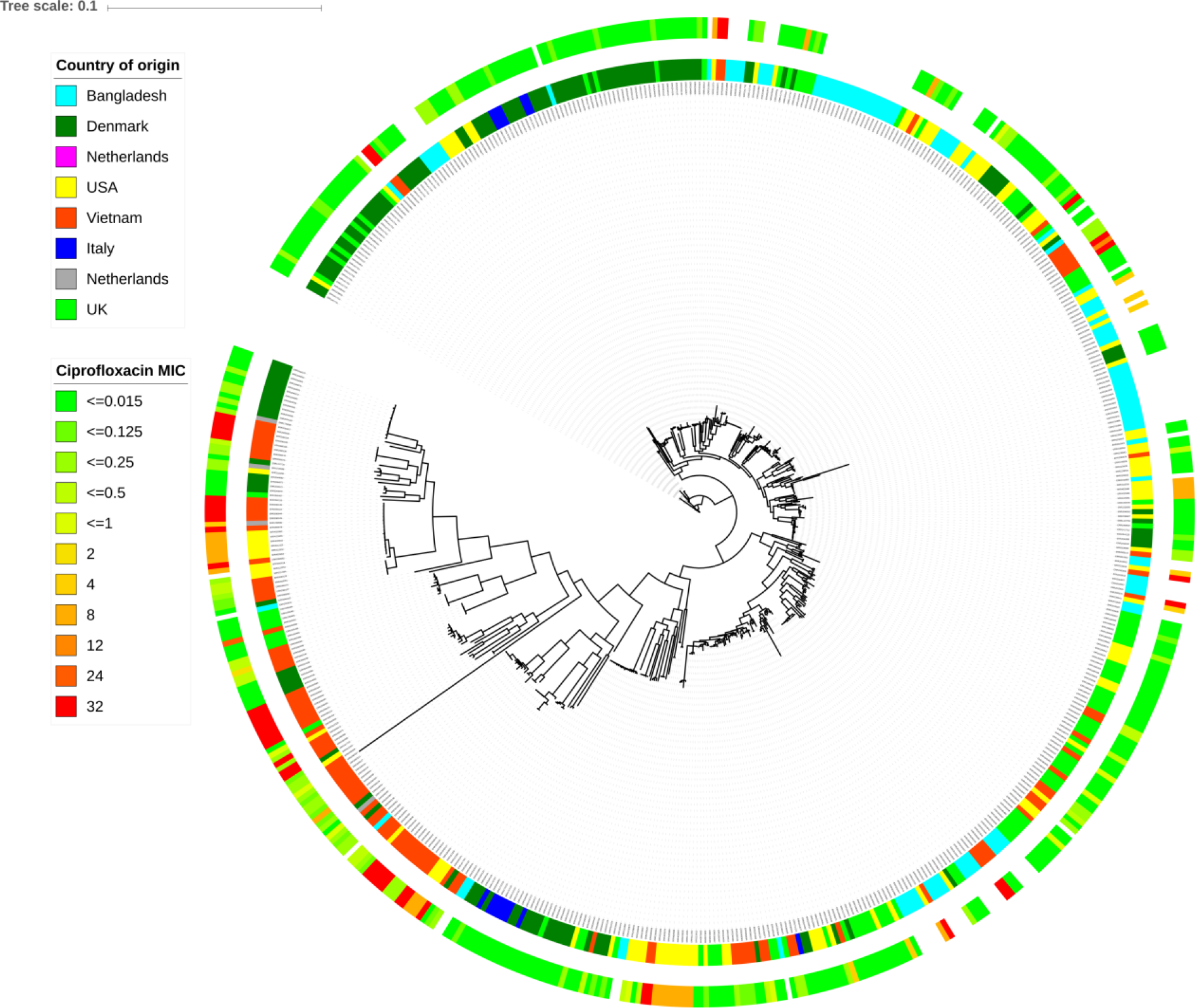
Maximum likelihood tree based on the alignment of concatenated core genes of 572 *E. coli* genomes (5 genomes failed quality control or assembly). Country of origin (inner circle) and ciprofloxacin minimum inhibitory concentration (outer circle) are indicated for each isolate. Zone diameters values for antibiotic susceptibility were not included in this analysis (isolates from Bangladesh).

## Discussion

We have built the AMR data hub with aim to provide a system for public health, food and veterinary institutes, clinical laboratories and researchers to share their genomic and related AST data. It can be used for standardized open-access data sharing, for example for published data, thus creating an ever-growing source of AST metadata available to researchers worldwide. The large volume of data made available will make it easier to use advanced statistical methods such as machine-learning to predict AMR phenotypes from genomic data and discover new AMR determinants.

It has been recently underlined that application of the Nagoya Protocol, which regulates material and data sharing, to genetic information might threaten the timely sharing of data in times of public health emergencies ^14^. In allowing the organisation and sharing of linked genomics and AST data, the AMR data hub promotes openness and accessibility for these important data types while at the same time meeting the privacy concerns for pre-publication data. Considering the exponential rise in the number of bacterial genomes available, and the threat to modern medicine represented by the rise of AMR, the establishment of the AMR data hub represents a timely effort to improve collaboration in this field.

The design of a standard data submission format benefited from the expertise of the COMPARE consortium, a group of international experts in bacterial genomics and AMR surveillance and research. It is designed to be as exhaustive, and at the same time as flexible as possible to ensure easy sharing of AST data. The database is hosted at the EMBL-EBI, ensuring its connection to the world’s largest online repository of bacterial genomes.

As members of the INSDC, EMBL-EBI and NCBI are part of a joined effort for standardization and sharing of genomic data. The NCBI can also host antibiograms in a similar format to that presented here, and efforts are currently ongoing to allow automated synchronization of content from both sides. This will greatly increase the flexibility and the reach of the AMR data hub. While NCBI data must be made public immediately upon deposition, EBI allows for pre-publication data to be kept private for a provider-defined period. Participating institutions can thus choose whether they want their data to be open-access immediately or whether they prefer sharing it with selected members of a consortium before public release.

The view is that the AMR data hub will soon become an essential resource for functional genomic studies of AMR. By encouraging data providers from different fields and geographical origin to share their data, this collection can greatly improve our ability to answer questions related to the current AMR crisis.

## Supporting information

Supplementary metarial and code

## Acknowledgements

This study was supported by the COMPARE Consortium, which has received funding from the European Union’s Horizon 2020 research and innovation programme under grant agreement No. 643476.

Contributing institutions for AST data: Amsterdam UMC^1^, APHA^11^, DTU^2^, FM^7^, OUCRU^13^, SSH^12^, UNIBO^10^.

