## Supplementary metarial and code for "Accelerating surveillance and research of antimicrobial resistance – an online repository for sharing of antimicrobial susceptibility data associated with whole genome sequences"

### Supplementary material

#### Methods: phylogenetic analysis of the collection of *E. coli* genomes

A list of the available genomes in the AMR data hub linked with antibiograms was generated using the Notebooks and the ENA Pathogens application programming interface (API) as described in the manuscript. The fastq sequence files were retrieved by querying the API using a password with command (code: 1. See below for code used).

The quality of the raw sequence reads was checked using fastqc (code: 2) (http://www.bioinformatics.babraham.ac.uk/projects/fastqc/). The species was verified to be *E. coli* for all isolates using KmerFinder 2.0 [9-10]: 4 isolates were discarded on suspicion of high contamination by non-E. coli reads. Reads were trimmed for quality and presence of linker sequences using Trimmomatic V0.33 [1] (code: 3). *De-novo* genome assembly was performed with SPAdes 3.9 [2] for Illumina short reads (code: 4) and with Canu v1.3 for PacBio long reads [3] (code: 5). Contigs of less than 500 bp long were removed from the genomes to improve the overall quality of the assembly using the perl script (removesmall.pl) written by Dr. Tamer Mansour and available at: https://github.com/drtamermansour/p_asteroides/blob/master/scripts/removesmalls.pl (code: 6) The quality of the assemblies was checked using quast [4] (code: 7). Identification of open reading frames (ORFs) and gene contents in the assembled genomes was performed using Prokka v1.11 [5] (code: 8). Core genome alignment was performed with Roary v3.6.8 [6] (code: 9).

The core gene alignment was used as input for calculation of distances and tree building using RAxML v8.1.6 [7] (code: 10).The phylogenetic tree and accompanying metadata was visualized using iTOL v3.3.1 (http://itol.embl.de/) [8].

All sequence manipulations described here were performed on the LISA Research Capacity Computing Services (https://userinfo.surfsara.nl/systems/lisa), using computation nodes equipped with E5-2650 v2 processors (16 cores), 64 Gb of RAM and running Debian Linux.

#### Code

1:

```
wget -nc --password [password] ftp://dcc_schubert@ftp.dcc-private.ebi.ac.uk/data/fastq/ERR204/XXX/[ACCESSION]/[ACCESSION].fastq.gz
```

2: (example for Illumina paired-end reads)

```
fastqc ACCESSION_R1.fastq.gz, ACCESSION_R2.fastq.gz -o fastqc_output
```

3: (example for Illumina paired-end reads)

```
java -jar trimmomatic-0.38.jar PE ACCESSION_R1.fastq.gz ACCESSION_R2.fastq.gz ACCESSION_R1_trimmed.fastq.gz ACCESSION_R1_failed.fastq.gz ACCESSION_R2_trimmed.fastq.gz ACCESSION_R2_failed.fastq.gz ILLUMINACLIP:./adapters/TruSeq3-PE-2.fa:2:30:10 SLIDINGWINDOW:4:15 LEADING:3 TRAILING:3 MINLEN:36
```

4:

```
spades.py -1 ACCESSION_R1_trimmed.fastq.gz -2 ACCESSION_R2_trimmed.fastq.gz -o spades_output_ACCESSION --careful
```

5:

```
canu -assemble -d ACCESSION_canu_out genomeSize=5m -p ACCESSION - pacbio-corrected ACCESSION.fastq.gz
```

6:

```
removesmalls.pl 500 spades_output_ACCESSION/ACCESSION_scaffolds.fasta > spades_output_ACCESSION/ACCESSION_scaffolds_cleaned.fasta
```

7:

```
python /quast-4.4/quast.py -s -o quast_output /genomes_cleaned/*.fasta
```

8:

```
prokka --outdir /prokka/ACCESSION --prefix ACCESSION /genomes_cleaned/ACCESSION_scaffolds_cleaned.fasta
```

9:

```
roary -p 15 -g 70000 -f /roary_output/ -e -n -v /prokka/*.gff
```

10:

```
raxmlHPC-PTHREADS-SSE3 -T 16 -m GTRGAMMA -p 31766040 -f a -x 7417 -N autoMRE -s /roary_output/core_gene_alignment.aln -n datahub_phylogeny -w /raxml_output/
```
